# Novel allosteric mechanism of p53 activation by small molecules for targeted anticancer therapy

**DOI:** 10.1101/384248

**Authors:** Joanna Zawacka-Pankau, Vera V. Grinkevich, Mikhail Burmakin, Aparna Vema, Karin Fawkner, Natalia Issaeva, Virginia Andreotti, Eleanor R. Dickinson, Elisabeth Hedström, Clemens Spinnler, Alberto Inga, Lars-Gunnar Larsson, Anders Karlén, Olga Tarasova, Vladimir Poroikov, Sergey Lavrenov, Maria Preobrazhenskaya, Perdita E. Barran, Andrei L. Okorokov, Galina Selivanova

## Abstract

Given the immense significance of p53 restoration for anti-cancer therapy, elucidation of the mechanisms of action of p53-activating molecules is of the utmost importance. Here we report a discovery of novel allosteric modulation of p53 by small molecules, which is an unexpected turn in the p53 story. We identified a structural element involved in p53 regulation, whose targeting by RITA, PpIX and licofelone block the binding of p53 inhibitors, MDM2 and MDMX. Deletion and mutation analysis followed by molecular modeling, identified the key p53 residues S33 and S37 targeted by RITA and PpIX. We propose that the binding of small molecules to the identified site induces a conformational trap preventing p53 from the interaction with MDM2 and MDMX. These results point to a high potential of allosteric activators. Our study provides the basis for the development of therapeutics with a novel mechanism of action, thus extending the p53 pharmacological potential.

## Introduction

Half of the human tumors express inactive mutant p53, while wild-type p53 is rendered functionally inert in cancer mainly due to the deregulated E3 ubiquitin ligase MDM2 and its homolog MDMX, which together potently inhibit p53 (Vousden et al. 2009). Several molecules targeting the p53-binding pocket of MDM2, such as nutlin and MI compounds, or the inhibitors of E3 ubiquitin ligase activity of MDM2, have been shown to induce p53-dependent growth suppression (Vassilev et al. 2004; Shangary et al. 2008; Yang et al., 2005) and some are currently tested in clinical trials (Hoe et al. 2014). However, MI compounds and nutlins do not inhibit another p53 inhibitor – MDMX, which makes them less efficient in tumors overexpressing MDMX protein (Toledo et al. 2007).

We have identified a small molecule RITA in a cell-based screen for the p53 reactivating compounds (Issaeva et al. 2004). RITA restores wild-type p53 in tumor cells by preventing p53/MDM2 interaction (Issaeva et al. 2004; Enge et al. 2009; Grinkevich et al. 2009). Next, we have found that protoporphyrin IX (PpIX), a metabolite of aminolevulinic acid, a pro-drug applied in photodynamic therapy of cancer, activates p53 by inhibition of p53/MDM2 complex (Zawacka-Pankau et al. 2007). In contrast to nutlin, RITA does not target MDM2 but binds to the p53 N-terminus (Issaeva et al. 2004). However, how the binding to p53 affects p53/MDM2 complex remains unclear.

In the present study, we applied state-of-the-art molecular and cell biology approaches and molecular modeling to map the region within the p53 N-terminus targeted by small molecules and to address the mechanism of their action. We found that RITA targeted p53 outside of the MDM2-binding locus and identified the key structural elements in the RITA molecule along with contact residues in p53, which are critical for the interaction. We found that the binding of RITA promotes a compact conformation of partially unstructured N-terminus, which inhibits the interaction with MDM2 and MDMX. Further, another p53 activator PpIX acts through a similar mechanism. Based on our results, we propose a model of a new allosteric mechanism of p53 activation. Using our model and chemoinformatic approaches, we have identified licofelone, a dual COX/LOX inhibitor, which blocks p53/MDM2 interaction via the mechanism that we uncovered.

## Results

### RITA selectively interacts with p53 in cancer cells

Our previous findings indicate that RITA interacts with the N-terminal region of p53 *in vitro* (Issaeva et al. 2004). To test whether RITA targets p53 in a cellular context, we analyzed [^14^C]-RITA complexes with proteins formed in HCT 116 colon carcinoma cells carrying wild-type p53 and in their p53-null counterparts (HCT 116 *TPp53-/-*). To visualize the complexes, we electrophoretically separated them and detected the position of RITA and p53 by autoradiography and Western blot, respectively. Under mild denaturing conditions (snap boiling in the loading buffer), [^14^C]-RITA migrated with the electrophoretic front in the lysates of HCT 116 *TPp53-/-* (Figure 1A), whereas in the lysates of HCT 116 cells its migration was shifted, indicating the formation of complexes. The position of the major band coincided with that of p53. Further, the immunodepletion of p53 from the lysates (Figure 1A) significantly decreased its intensity supporting the notion that it represents p53/RITA complex.

**Figure 1.**
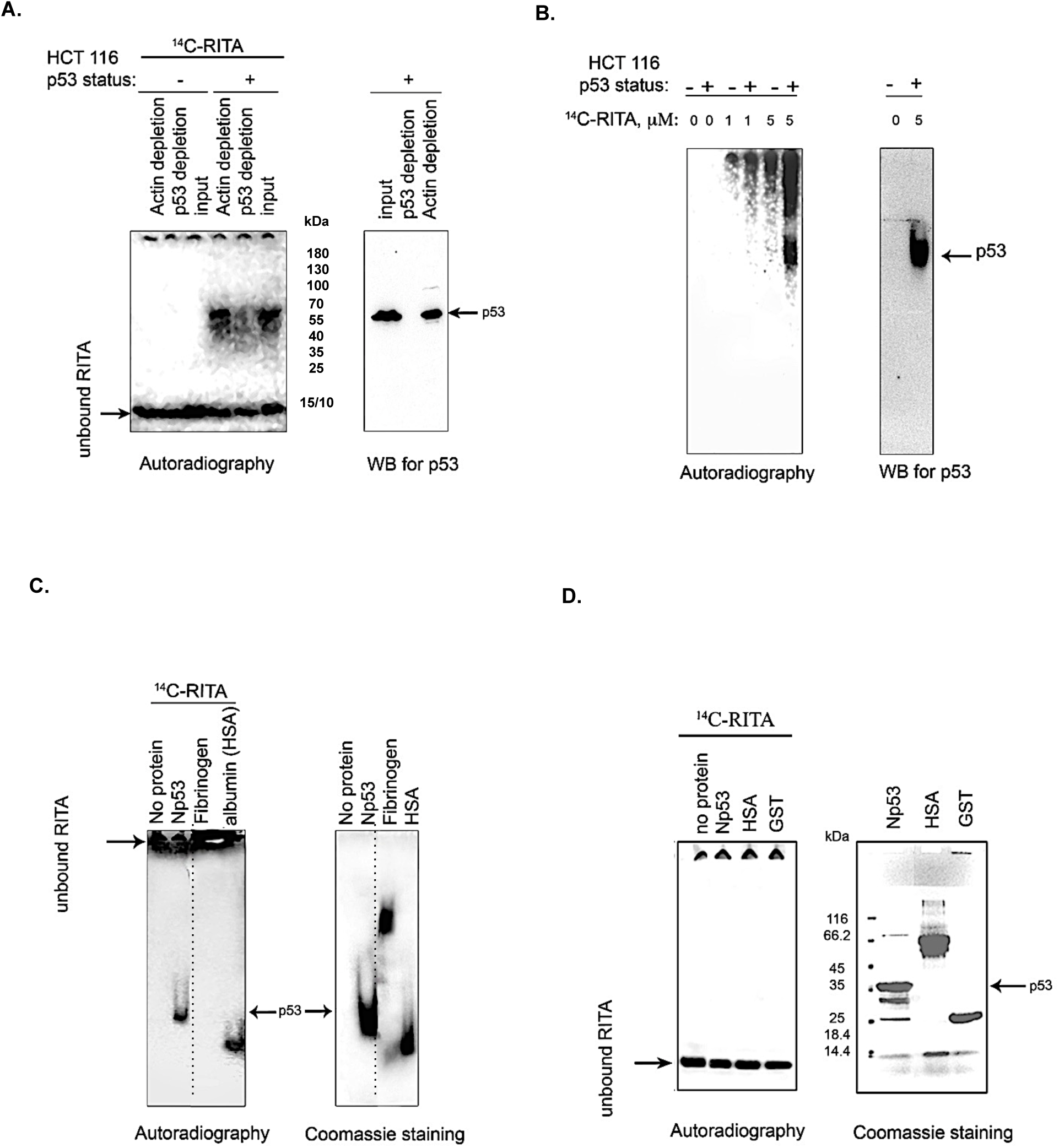
RITA binds p53 in cells and *in vitro*. **A.** [^14^C]-RITA/protein complexes were detected in lysates of HCT 116 or HCT 116 *TP53/-* cells treated with 5 μM [^14^C]-RITA for 12h upon separation in 10% SDS-PAGE under mild denaturing conditions. The position of RITA in the gel was visualized by autoradiography. p53 was detected by immunoblotting with the DO-1 antibody (right panel). Shown is a representative data of three independent experiments. **B.** A small-molecule band shift assay was used to detect [^14^C]-RITA/protein complexes in lysates of HCT116 or HCT116*TP53-/-* cells treated with [^14^C]-RITA, and separated in gradient polyacrylamide gel under native conditions. [^14^C]-RITA and p53 were detected as in **A**. **C.** [^14^C]-RITA interaction with recombinant proteins, GST-Np53(2-65), fibrinogen and human serum albumin (HSA) was detected by band shift assay using 2:1 molar excess of RITA. The dotted line indicates where the gel was cut. **D.** Upon standard SDS electrophoresis [^14^C]-RITA/protein complexes were disrupted (n=3).

Next, we developed a small-molecule band shift assay, analogous to the commonly used band shift assay for detection of protein/DNA binding. We separated [^14^C]-RITA/cellular proteins complexes by native electrophoresis and detected [^14^C]-RITA by autoradiography (Figure 1B). The major band of RITA/protein complex in HCT 116 cells treated with 5 μM coincided with that of p53 (Figure 1B). The absence of a similar band in the p53-null cells (at 5 μM RITA) (Figure 1B) indicates that it represents RITA bound to p53. We detected weaker RITA/protein complexes in HCT 116 p53-/-. We are not ruling out the possibility that RITA might interact with other proteins in cells lacking p53. One of the possible binding partners might be the p53 protein family member, p73 tumor suppressor since it bears high structural homology with the p53 N-terminus and is regulated by MDM2 and MDMX proteins. Taken together, our data provide evidence for the selective interaction of RITA with p53 in cancer cells.

### RITA interacts with the N-terminus of p53

Next, we analyzed the interaction of RITA with the recombinant p53 N-terminus employing our small-molecule band shift assay. Upon incubation, ^14^C-RITA formed a complex with Glutathione-S-transferase (GST)-fusion p53 N-terminus (Np53) (2-65), as manifested by the co-migration with Np53 (Figure 1C) but only weakly interacted with GST-tag (Figure 2B). In contrast, RITA did not associate with the human fibrinogen (Figure 1C), suggesting a selective interaction with p53. Human serum albumin (HSA), a known carrier of various drugs in the blood (Koehler et al. 2002), was used as the binding control (Figure 1C). Under denaturating conditions [^14^C]-RITA/protein complexes were disrupted (Figure 1D), suggesting that this interaction is reversible.

**Figure 2.**
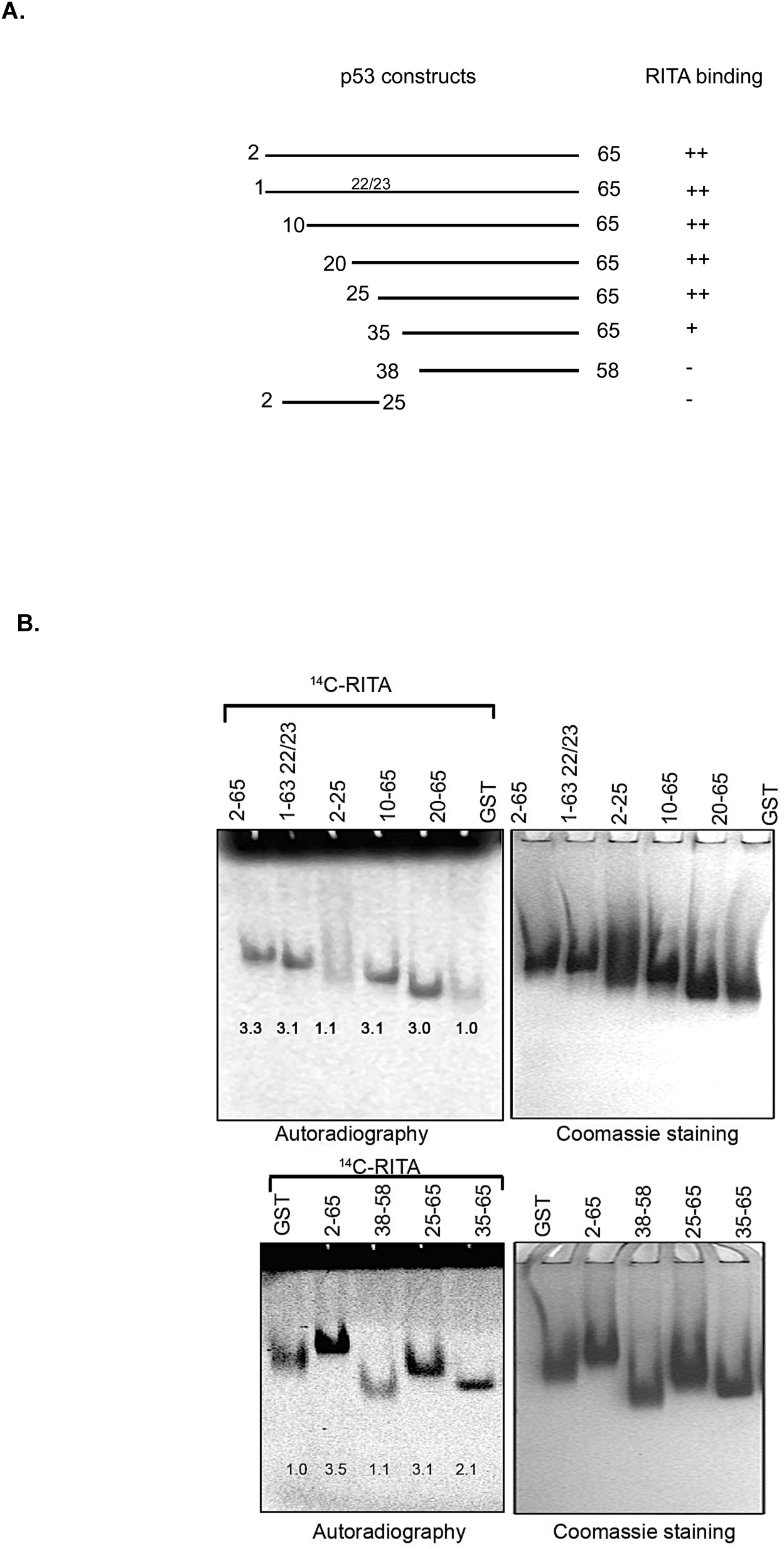
Mapping of RITA binding site within the p53 N-terminus. **A.** Scheme depicting the series of deletion mutants of GST-Np53 used in this study. **B.** The binding of [^14^C]-RITA to p53 N terminus deletion mutants was analyzed in band shift assay as in Figure 1C. Band density was measured using ImageJ software and normalized to GST-tag signal. This is a representative data of three independent runs.

Non-labeled RITA readily competed out the [^14^C]-RITA from the complex with Np53 at a low molecular excess, 1:1 or 1:2.5 (**Figure S1A**). However, it did not effectively compete with the [^14^C]-RITA/HSA complex (**Figure S1B**) suggesting a different mode of interaction.

### Mapping RITA binding site using deletion and point mutants

To identify key p53 residues involved in the binding to RITA, we applied deletion mutagenesis approach to generate a series of p53 deletion mutants and assessed their interaction with RITA (Figure 2A).

Deletion of the first 25 residues containing the MDM2 binding site or mutations in residues 22/23 required for the interaction with MDM2 did not change the binding of p53 to RITA (Figure 2B). These results, as well as a weak, if any, interaction with Np53(2-25) protein (Figure 2B, upper panel) argue against the binding of RITA within the MDM2 site of p53.

Notably, Np53(38-58) GST fusion protein did not interact with RITA either (Figure 2B, lower panel). Together, our results indicate that RITA target sequence is located between residues 25-38 (Figure 2A and B). Further analysis revealed that Np53(35-65) interacted with RITA approximately 50% less efficiently than Np53(2-65) (Figure 2B). Therefore, we concluded that RITA targets residues located in the proximity to leucine 35.

### Molecular modeling of RITA/p53 complex

The X-ray crystallographic analysis of the p53-MDM2 complex structure shows that the N-terminal p53 region binds the MDM2 hydrophobic groove in the a-helical form (Kussie et al., 1996) and the formation of this complex represents an important disordered-to-ordered transition (Lee et al., 2000). The N-terminal region of p53 is largely disordered and highly flexible and forms an amphipathic helical structure in the vicinity to MDM2 (Uversky, 2016).

Thus, based on the limited, available information on the structural organization of the p53 N-terminus (Okorokov et al., 2006; Lowry et al., 2008; Espinoza-Fonesca, 2009) and our deletion mutagenesis, we performed Monte Carlo conformational search to explore the possible binding modes of RITA to the p53 N-terminus (MacroModel, 2008). The MCMM-LMOD search on the RITA-p53 complex found 3492 low energy binding modes within 5 kcal/mol above the global minimum. Among these, the tenth lowest energy binding mode, 2.1 kcal/mol above the global minimum, appeared reasonable concerning the placement and orientation of RITA molecule.

This model implies that the binding of RITA involves the formation of hydrogen bonds between its terminal hydroxyl groups and serine 33 and serine 37 of p53, as well as hydrophobic interactions with proline 34 and 36 via one of its thiophene and the furan rings (Figure 3A and B and **Supplemental video 1**). Hydrogen bonds and hydrophobic interactions between RITA and the p53 SPLPS amino acid sequence increase the already limited flexibility of this region (Figure 3A and B).

**Figure 3.**
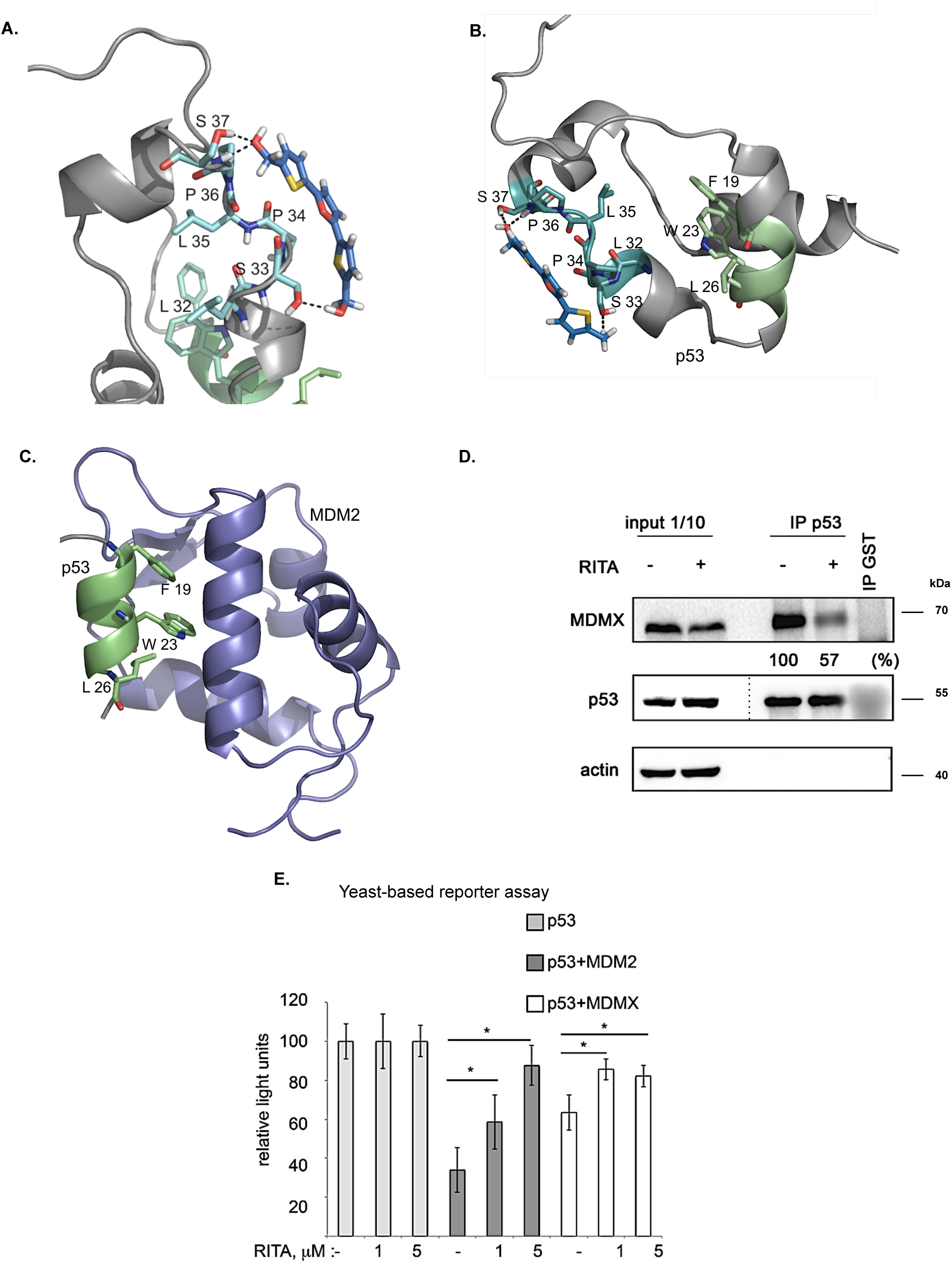
Molecular model of the p53/RITA complex and inhibition of the p53/MDMX interaction by RITA. **A.** Binding of RITA to SPLPS sequence (cyan) of p53 involves interaction with S33 and S37 via terminal hydroxyl groups of RITA and hydrophobic interactions with P34 and P36. Hydrogen bonds are highlighted in black dotted lines. The orientation of MDM2-binding helix of p53 (*lime*) is different upon p53 binding to RITA (blue) **B.** and MDM2 (*purple*) (pdb: 1YCQ) (**c**). Side chains of residues (F19, W23, and L26) involved in MDM2 binding are shown in (**B.**, **C**.). Atom type coloring; oxygen (*red*), nitrogens (*blue*), and sulfurs (*yellow*). See also **Movie S1**. **D.** In line with the model prediction, RITA-mediated p53 conformational change results in the inhibition of p53/MDMX binding in HCT 116 cells as assessed by co-immunoprecipitation. This is representative data of three independent experiments. The dotted line indicates the site where the membrane was cut. **E.** RITA rescues the p53 transcriptional activity from inhibition by MDM2 or MDMX as assessed in a yeast-based functional assay. The average light units relative to the transactivation activity of p53 alone and the standard errors of at least five biological repeats are presented. The *t*-student test was performed for statistical analysis with p = 0.05.

Molecular dynamics simulations suggest that leucine-rich hydrophobic clusters within residues 19-26 and 32-37 stabilize the folding and formation of α-helixes in the N-terminus (Espinoza-Fonesca et al. 2009). According to this study, MDM2-contacting residues F19, W23 and L26 within the α-helix of p53 (residues 16-26) are facing inwards and are tacked inside, stabilized by the formation of hydrophobic leucine clusters, while more hydrophilic residues of the α-helix are exposed to the solvent. This is supported by the tryptophan fluorescence assay, which demonstrated that W23 is shielded from the solvent (Kar et al., 2002). On the other hand, the X-ray structure of the MDM2-p53 peptide complex (1YCQ.pdb) shows that MDM2-contacting residues are facing out (Figure 3C). This indicates that the binding to MDM2 requires a partial unwinding of the α-helix to flex out F19, W23, and L26, as illustrated in Figure 3B and 3C. Interestingly, a study by Lum and colleagues showed a slowed kinetics of loop closure in segments 13-23, 23-31, 31-53 and 53-60 of the p53 N-terminus induced by phosphorylation of S33, S46, and T81. This indicates that local electrostatic changes in p53 N-terminus induced by phosphorylation can be transmitted to remote sites through transient interaction networks in the disordered domain (Lum et al., 2012).

In line with this observation, our model indicates that RITA, by increasing the rigidity of the proline-containing SPLPS motif, induces a conformational trap in a remote MDM2 binding site. Next, we propose that constraints imposed by RITA prevent solvent exposure of F19, W23, and L26 residues, thus counteracting the p53/MDM2 interaction (Figure 3B and 3C).

Conformational change induced by RITA is expected to impinge on other protein interactions involving the p53 N-terminus. The binding of p53 to the MDM2 homolog, MDMX requires the formation of an α-helix as well as exposure of the same p53 residues, as facilitated by MDM2. We, therefore, reasoned that the conformational change induced by RITA might, also, abrogate the binding of MDMX as well.

### RITA inhibits p53/MDMX interaction in cells and in vitro

To assess whether RITA could inhibit p53/MDMX complex, we treated HCT 116 colon cancer cells with RITA and performed co-immunoprecipitation. Our data indicated that RITA reduced the amount of MDMX bound to p53 by 43% (Figure 3D).

To further elucidate the ability of RITA to inhibit the p53/MDM2 and p53/MDMX interactions, we employed a yeast-based assay, which measures p53 transcriptional functionality using as readout the activity of a p53-dependent luciferase reporter. Since MDM2 does not degrade p53 in yeast cells, the inhibitory effect of MDM2 in this system is ascribed to the direct interaction with p53 and consequent inhibition of p53-dependent transcription ^1^. Co-transfection of MDM2 inhibited the transcription activity of p53 (Figure 3E), whereas RITA rescued wtp53-mediated transactivation of the reporter. In addition, RITA protected p53 from inhibition by MDMX (Figure 3E).

Taken together, our results demonstrated that the allosteric effects exerted by RITA result in the inhibition of both p53/MDM2 and p53/MDMX interactions.

### Terminal hydroxyl groups of RITA are crucial for RITA/p53 interaction

Our model implies that the central furan ring of RITA is not relevant for the binding with p53. Indeed, an analog of RITA with the substitution of furan oxygen atom to sulfur (LCTA-2081, compound 2, see **ST1** for structure) had comparable p53-dependent activity in HCT 116 cells as assessed by the viability assay (Figure 4A). Further analysis of RITA analogs (**ST1**) let us conclude that the presence of three rings is required for its p53-dependent biological activity.

**Figure 4.**
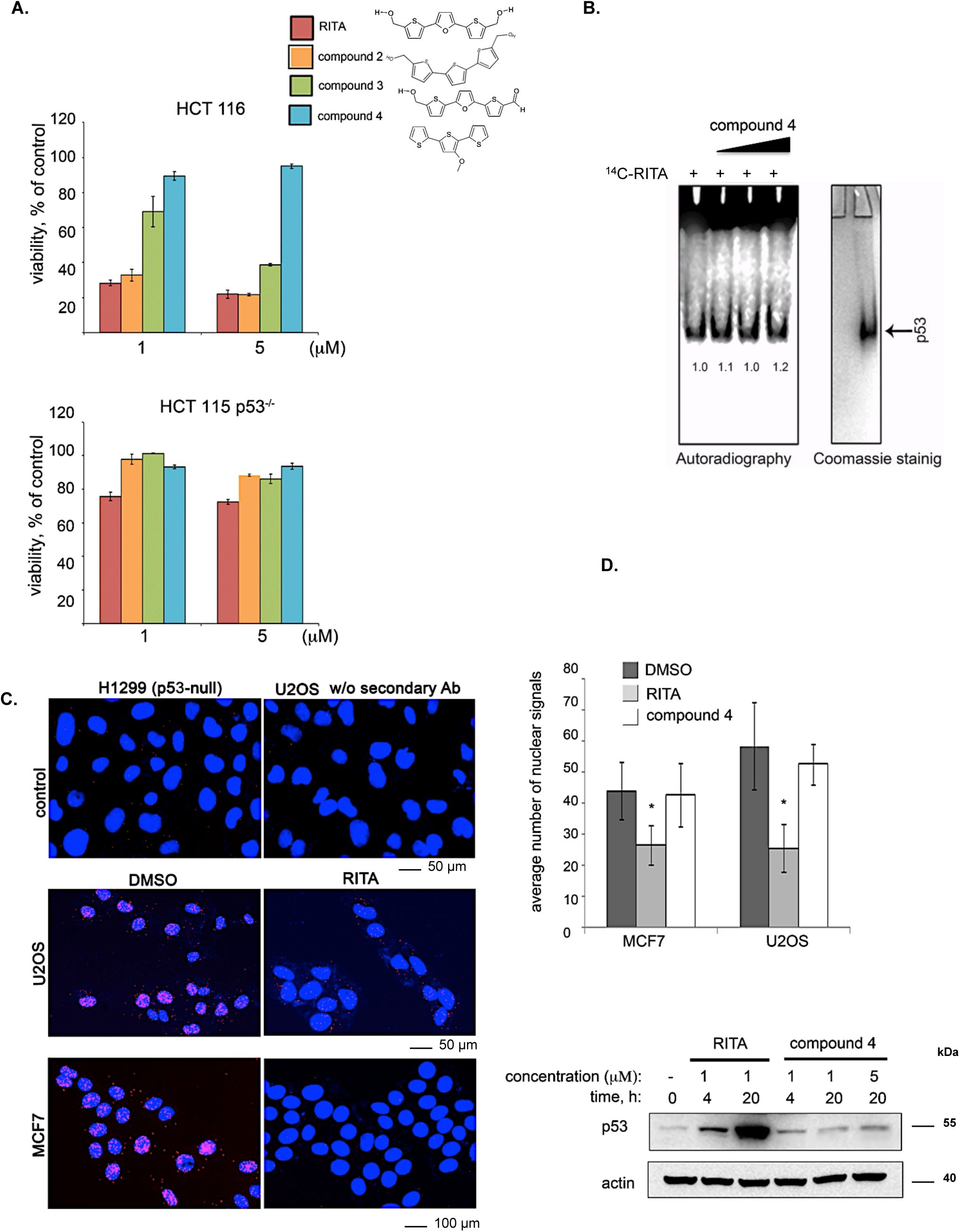
Two terminal hydroxyl groups of RITA are crucial for the binding to p53 and disruption of p53/MDM2 interaction. **A.** RITA analog NSC-650973 (compound 4) lacking two hydroxyl groups did not suppress the growth of HCT 116 cancer cells, unlike LCTA-2081 (compound 2) analog with substituted O atom in furan ring (for structure refer to **Supplementary Table 1**), which retained full biological activity. NSC-672170 (compound 3) analog with one hydroxyl group substituted to ketone retained more than 60% of RITA biological activity. **B.** Biologically inactive RITA analog compound 4 (40, 80 and 100 μM) did not compete for the binding to GST-Np53 with [^14^C] -RITA. **C.** Visualization of p53/MDM2 complexes in MCF7 and U2OS cells treated or non-treated with RITA for 6 hrs using *in situ* Proximity Ligation Assay (isPLA) (red fluorescent dots). The p53-null H1299 and U2OS cells stained without secondary antibody were used as the assay controls. **D.** isPLA demonstrated the decrease in the average number of nuclear signals upon treatment with RITA, but not with its derivative NSC-650973 (upper panel). The normality was assessed with Shapiro-Wilk’s test. p< 0.05 values were considered statistically significant. RITA, but not compound 4 induced p53 accumulation in HCT116 cells, as detected by immunoblotting (lower panel).

Our molecular modeling predicts that one or two terminal hydroxyl groups are key for the interaction with p53. This prediction was supported by the loss of biological activity of RITA analog NSC-650973 (compound 4, **ST1**), lacking both hydroxyl groups (Figure 4A). Further, this compound did not compete with [^14^C]-RITA for the binding to p53 *in vitro* (Figure 4B), when the samples of GST-dNp53 (20 μM) and [^14^C]-RITA (40 μM) were incubated with the increasing concentrations (40, 80 and 100 μM) of the inactive analog 650973-N indicating that it does not interact with p53. Compound 4 contains additional methoxy group that adds ups to the change of the structure when compared to RITA. However, further experiments highlight the crucial relevance of RITA hydroxyl groups and Np53 serine residues for RITA/Np53 complexes.

Next, we assessed the inhibition of p53/MDM2 interaction by RITA and compound 4 in cells using *in situ* proximity ligation assay (isPLA) (Figure 4C) (Söderberg et al., 2006; Castell et al., 2018). isPLA allows for detection of the interaction between the proteins using specific antibodies tagged to oligos. Briefly, the red fluorescence signals localized mostly in the nucleus indicate p53/MDM2 interactions. Treatment of MCF7 or U2OS cells with RITA significantly decreased the average number of p53/MDM2 isPLA nuclei signals (from 44 to 26.5 in MCF7 cells and from 58.32 to 25.52 in U2OS cells when compared with DMSO). Unlike RITA, compound 4 did not decrease the average number of isPLA nuclei signals, indicating that it does not inhibit p53/MDM2 interaction (Figure 4D, upper panel). In line with these data, compound 4 did not induce p53 accumulation (Figure 4D, lower panel). Notably, compound 3, lacking one hydroxyl group (**ST1**, **Figure S2B**) was more efficient in suppressing the growth of HCT 116 cells than compound 4 (NSC-650973) but still less potent than RITA (Figure 4A and Issaeva et al., 2004). Thus, we conclude that in agreement with our model, both terminal hydroxyl groups of RITA and three thiofuran rings are required for the efficient binding to p53. The ability to bind p53 correlates with the prevention of p53/MDM2 binding, induction of p53 and p53-dependent growth suppression.

### Serine 33 and serine 37 are critical for RITA/p53 interaction, p53 stabilization, and transcription activity

To further validate our model, which predicted the crucial role of serines 33 and 37 for RITA interaction, we mutated serine 33 (S33) to alanine, alone or in combination with serine 37 (S37), and assessed the binding of RITA to Np53(S33A) and Np53(S33A/S37A) (referred to as p53 (33/37) peptide) using band-shift assay.

In line with our model, the interaction of both mutant proteins with RITA was significantly decreased (Figure 5A), indicating the key role of these residues in binding to RITA. The difference in the intensity of the radioactive signal detected for the GST-tag is due to the usage of different phosphor screens and different batches of purified proteins. The variations in the signal intensities do not affect the general conclusion that RITA interacts strongly with wt Np53 and displays only background binding to GST-tag.

**Figure 5.**
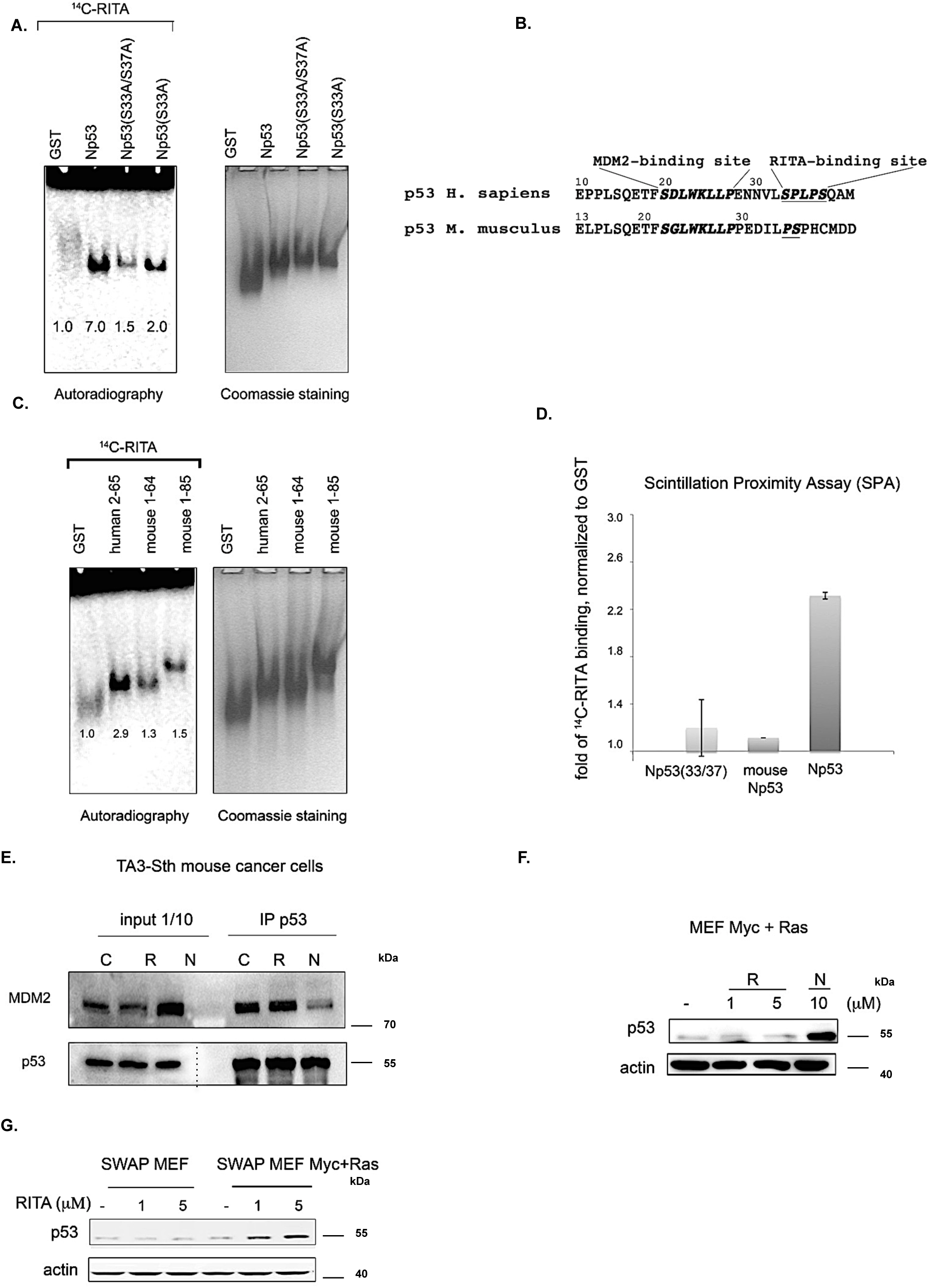
Serine 33 and serine 37 are crucial for the efficient binding of RITA to the p53 N-terminus. **A.** Assessment of [^14^C]-RITA interaction with Np53 proteins carrying alanine substitutions of S33 or S33/S37 using band shift assay as in Figure 1C. Bands’ density was quantified using ImageJ software and normalized to GST-tag. **B.** Alignment of mouse and human p53 N-termini, highlighting the site for MDM2 interaction and RITA-binding motif. **C. D.** Inefficient binding of RITA to GST-fusion mouse p53 proteins, spanning residues 1-64 and 1-85 as detected by band shift assay and scintillation proximity assay (SPA). SPA assay reveals inefficient binding of RITA to Np53(33/37). Bands’ density was quantified using ImageJ software and normalized to GST-tag (**C**). **E.** Co-immunoprecipitation experiment demonstrated that RITA did not prevent MDM2/p53 interaction in TA3-Sth mouse cancer cells, unlike nutlin. C - control untreated sample, R - RITA-treated and N - nutlin-treated samples; the dotted line represents different exposure time of this part of the membrane. **F.** Mouse p53 in Myc- and Ras-transformed MEF’s was not induced by RITA (R) in contrast to nutlin (N). **G.** RITA efficiently induced the level of human p53 in SWAP MEF’s transfected with Ras and c-Myc as detected by immunoblot.

Comparing to human p53, mouse p53 lacks residues corresponding to serine 33 and proline 34, a residue, which is responsible for the hydrophobic interaction with RITA furan ring. Thus, to further highlight the relevance of the SPLPS motif in the binding of RITA to Np53, we decided to test whether RITA can bind mouse Np53 (Figure 5B). We detected only a 0.3 and 0.5 increase above the GST-tag in the radioactive signal of RITA bound to mouse Np53(1-64) and Np53(1-85) respectively. The increase in the radioactive signal above the GST-tag for human Np53 was 1.9 (Figure 5C), suggesting that the presence of S33 and P34 residues is important for RITA binding to p53. Further, in line with other assays, Scintillation Proximity Assay (SPA), which detects the excitation of protein-coated beads by radioactively labeled RITA only when in very close proximity (24), revealed that the binding of ^14^C-RITA to mouse p53 and Np(33/37) mutant is inefficient (Figure 5D).

In contrast to nutlin, which blocked the p53/MDM2 complex and induced p53 accumulation in mouse cells, RITA did not disrupt the mouse p53/MDM2 interaction and did not induce p53 in mouse tumor cells and mouse embryonic fibroblasts (MEFs) expressing Ras and c-Myc oncogenes (Figure 5E and 5F). Nutlin but not RITA activated p53 beta-gal reporter in T22 mouse fibroblasts (**Figure S3**). These data are consistent with our previous results demonstrating the absence of growth suppression by RITA in mouse tumor cell lines (Issaeva et al., 2004).

Notably, swapping mouse p53 to human p53 in mouse embryo fibroblasts (SWAP MEF) derived from transgenic mice expressing human p53 in mouse p53-null background (Dudgeon et al., 2006) restored the ability of RITA to induce p53. As shown in Figure 5G, RITA induced p53 in SWAP MEF’s expressing c-Myc and Ras. It did not affect SWAP cells without Ras and Myc overexpression, which is in line with our previous data suggesting that oncogene activation, is required for RITA-mediated induction of p53 (Issaeva et al., 2004; Grinkevich et al., 2009). Taken together, these data suggest that S33 within SPLPS motif is required for RITA/p53 binding.

To further validate the role of Ser 33 and 37 we compared the ability of RITA to rescue wtp53 and S33A/S37A mutant (referred to as p53 (33/37) from MDM2 using yeast-based reporter assay. Both 10 μM nutlin and 1 μM RITA relieved p53-mediated transactivation from MDM2 inhibition (Figure 6A), whereas their effects on the activity of p53(33/37) were different. Nutlin protected both wt and p53(33/37) from inhibition by MDM2 equally well (*t*-student; p<0.05), but RITA had a significantly weaker effect on the p53(33/37) (Figure 6A). Due to short treatment, significant binding of ^14^C-RITA to p53 protein in cells was detected for a higher dose of the compound at which p53 accumulates rapidly in cancer cells. Potent inhibition of p53/MDM2 interactions at 1 μM RITA in yeast-based assay correlates well with the inhibition of p53/MDM2 complex in cancer cells (Issaeva et al., 2002) and indicates that disruption of p53/MDM2 and p53/MDMX interactions are needed for p53 stabilization and accumulation in cancer cells. These data lend further support to the notion that S33 and S37 play an important role in RITA-mediated inhibition of p53/MDM2 interaction.

**Figure 6.**
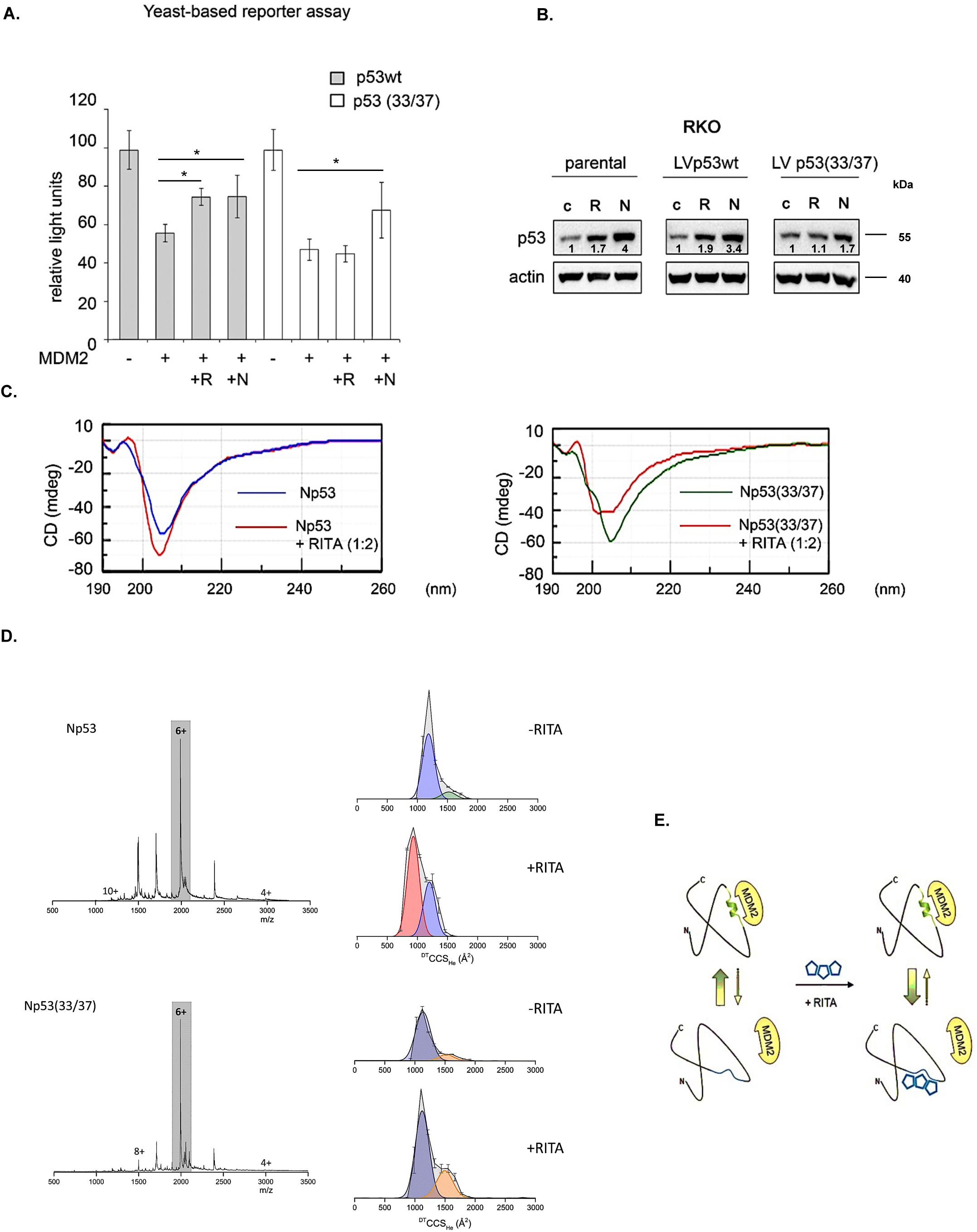
Serine 33 and serine 37 are required for RITA-mediated rescue of p53 from MDM2-dependent inhibition. **A.** The effect of the co-expression of MDM2 along with wt or 33/37 p53 upon the treatment with 1 μM RITA (R) or 10 μM nutlin (N) on a yeast-based luciferase reporter was assayed. The average light units relative to the transactivation activity of p53 alone and the standard errors of at least five biological repeats are presented. The *t*-student test was performed for statistical analysis with p*<*0.05. **B.** Induction of the p53 protein levels by 1 μM RITA (R) and 10 μM nutlin (N) in RKO colon cancer cells and their p53-null derivatives with reconstituted wt p53 and p53(33/37) as assessed by Western blotting. Band density was assessed using ImageJ software and normalized to non-treated controls. **C.** RITA promotes an increase in the secondary structure in wt Np53 (left) but not in Np53(33/37) (right) as detected by circular dichroism spectroscopy (CD). **D.** nESI mass spectra (left) and drift tube ion mobility mass spectrometry collision cross section distributions arising from arrival time distributions (right) for the [M+6H]^6+^ analyte of wt Np53 in the absence and presence of RITA (top panel, adapted from ^2^ and Np53(33/37) in the absence and presence of RITA (bottom panel). Coloured Gaussian curves depict conformational families. wtNp53 undergoes a compaction event resulting in the induction of a novel conformational family shown in red. Np53(33/37) conformational spread is unaffected by RITA induction. **E.** A scheme is illustrating the allosteric mechanism of RITA-mediated prevention of p53/MDM2 interaction. Binding of RITA shifts the balance towards p53 conformation with low affinity to MDM2.

Next, we addressed the question whether the same serine residues are important for the induction of p53 in human cells by RITA. We stably expressed S33/S37 p53 and wtp53 in colon carcinoma RKO *TP53-/-* cancer cells, in which both alleles of wtp53 were inactivated by homologous recombination (Sur et al., 2009). Nutlin induced the accumulation of wt and p53(33/37) with similar efficiency (Figure 6B and not shown). In contrast, the induction of the double serine mutant by RITA was impaired (Figure 6B and not shown).

Importantly, CD spectroscopy confirmed our previously published data (Dickinson et al. 2015) that RITA increases the content of the secondary structure in Np53 (Figure 6C, left panel). These data support our model that the binding of RITA induces a conformational change in p53 (Figure 6C, left panel). To elucidate the role of serine 33 and 37 in a RITA-mediated increase of the secondary structure in Np53, we incubated Np53(33/37) with the access of RITA (1:2 ratio) and performed CD analysis. As shown in Figure 6C, (right panel), RITA did not increase the secondary structure in Np53(33/37). In contrast, we observed the increase in the unstructured content after incubation with RITA. The change in Np53(33/37) upon RITA incubation remains to be elucidated.

Next, we applied ion mobility mass spectrometry (IM-MS) that allows studying the topology of proteins under various conditions (Jurneczko et al. 2013; Harvey et al. 2012). Briefly, we first incubated both wt and Np53(33/37) in the presence or absence of RITA. Proteins (50μM) were incubated with RITA in 1:2 ratio and analyzed by IM-MS (**Supplemental Experimental procedures**). The wtNp53 after incubation with RITA presents as ions of the form [M+zH]^z+^ where 4= *z* =10 with charge states 5= *z* =8 at a significant intensity (Figure 6D, upper panel). The mass spectra for Np53 without RITA, with RITA and the control spectra, show no mass shift, suggesting that RITA binding is lost during desolvation (**Figure S4A)**. Nonetheless, the conformation of Np53 in the presence of RITA was changed. The collision cross-section distributions in Figure 6D, show that in the absence of RITA, Np53 presents in two distinct conformational families centered at ∼1500 and ∼1750 Å^2^. Upon incubation with RITA the more extended conformer is lost, the conformer at ∼1500 Å^2^ remains present at a lowered intensity and a third conformational family centered on ∼1000 Å^2^ appeared, suggesting significant compaction of the protein. Control experiments confirmed that this conformer was present only after incubation with RITA (**Figure S4A**). This trend was observed for all sampled charge states (**Figure S5A**) with small variations in conformer intensity attributable to coulombic repulsion upon desolvation. Thus, RITA induces the generation of a unique compact conformer (or closely related conformational family) in wtNp53. This is in agreement with our previously published data showing that RITA promotes an allosteric change in p53 N terminus ^2^. In contrast, Np53(33/37) showed no gross conformational change after incubation with RITA (Figure 6D, bottom panel, and **Figure S4B**, **S5B**). Np53(33/37) presents in two conformations centered at ∼1100 and 1500 Å^2^ both in the absence and presence of RITA.

Taken together, these data provide compelling evidence that the allosteric activation of p53 by RITA requires residues S33 and S37.

### Identification of PpIX as another allosteric activator of p53

Next, we decided to assess whether the identified allosteric mechanism of p53 activation could be applied to other inhibitors of p53/MDM2 interaction. We have previously shown that small molecule protoporphyrin IX (PpIX), a photosensitizer applied in clinics, binds to the p53 N-terminus and disrupts p53/MDM2 complex (Zawacka-Pankau et al., 2007, Sznarkowska et al., 2010). Here, we tested if PpIX targets the same amino acid residues in p53 as RITA, using fluorescent-based small-molecule band shift assay. Fluorescent band shift assay indicates that substitution of serine 33 to alanine or double mutation in serine 33 and serine 37 decreases the binding of PpIX to the p53 N-terminus (**Figure S6**).

Next, we used a new technology called a Fluorescent-2 Hybrid (F2H^®^) Assay developed to study protein-protein interactions in live cells (Zolghard et al., 2008). Briefly, we employed tethering of MDM2 protein (LacI-GFP-Mdm2) at protein-protein interaction platform in the nucleus of U2OS cells and detected the complex of MDM2 with p53 (RFP-p53) using fluorescent-based imaging (**Figure S7A**). We detected a potent inhibition of p53/MDM2 interaction in U2OS cells by PpIX (61 ± 8%, *p*<0.01, n=6), which was comparable to positive control, nutlin (60 ± 5%, *p*<0.001, n=6).

Using the yeast-based reporter assay, we confirmed that PpIX rescues p53 transcriptional activity from both MDM2 and MDMX (**Figure S7B**).

Taken together, our findings implicate the conformational state of the SPLPS sequence distal from the MDM2-interacting residues as a key structural element regulating p53/MDM2 interaction as presented in the model in Figure 6E. This site could be modulated by small molecules such as RITA and PpIX.

### Identification of licofelone as an allosteric activator of p53

To identify new small molecules, which can prevent p53/MDM2 binding via the allosteric mechanism that we discovered, we performed an informational search using the Therapeutic Targets Database (TTD) (http://bidd.nus.edu.sg/group/cjttd/TTD_HOME.asp). We used the TTD internal Drug Similarity Search engine to estimate the Tanimoto coefficient (TC) of similarity for two molecules *A* and *B* (as described in **Supplemental Experimental procedures**). We identified several candidate compounds among which licofelone, a dual LOX/COX inhibitor, was the top compound with TC of 0.73 (**ST2**).

To select the candidate molecule for further studies, we next performed a function-related search of RITA and licofelone properties using computer program PASS, which predicts the biological activity spectra based on the structural formula of the compounds (Lagunin et al., 2010; Filimonov et al. 2014; 2018). The prediction analysis showed that 17 out 26 (65.4%) biological activities predicted for licofelone coincided with those predicted for RITA.

Thus, we selected licofelone for further validation in biological assays. We found that licofelone inhibited the growth of cancer cells (Figure 7A and B). The inhibition was p53-dependent. At higher doses, licofelone also suppressed the growth of HCT 116 p53-/- cells (Figure 7B). We speculate that the effect might be due to the activation of p53 protein family member, p73 protein. However, this yet needs to be unequivocally demonstrated. Next, licofelone efficiently competed for the binding to p53 in gel shift assay, indicating that it binds the p53 N-terminus (Figure 7C). In line with p53-dependent growth suppression, licofelone induced p53 accumulation and expression of p53 pro-apoptotic targets Puma and Noxa (Figure 7D). Pull down assay showed that licofelone disrupted p53/MDM2 interaction (Figure 7E), which supports its functional similarity to RITA.

**Figure 7.**
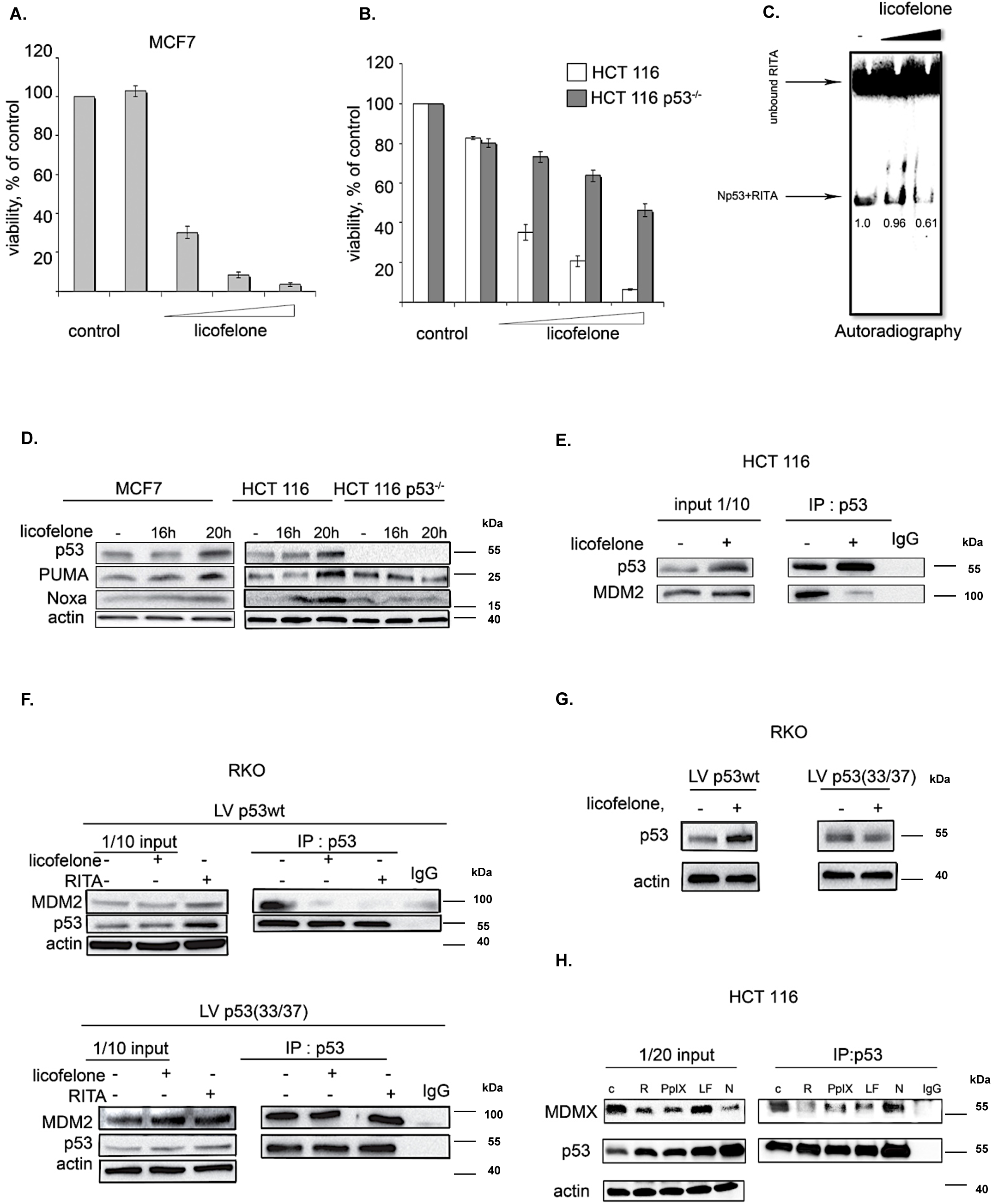
Licofelone activates p53 and disrupts p53/MDM2 and p53/MDMX interactions via targeting serine 33 and 37 residues. **A.** Dose-dependent inhibition of proliferation by licofelone (25, 50, 75 and 150 μM) in MCF7 cells as examined in WST assay. **B.** Licofelone (25, 50, 75 and 150 μM) inhibited the growth of human colon cancer cells in a p53-dependent manner. **C.** Licofelone competed for the binding to Np53 with ^14^C-RITA in small molecule band-shift assay. Bands’ density was measured using ImageJ and normalized to Np53 incubated with ^14^C-RITA alone. **D.** Licofelone induced p53 and its targets PUMA and Noxa in HCT 116 and MCF7. **E.** p53/MDM2 complex disruption in HCT 116 cells by licofelone as assessed in a co-immunoprecipitation assay. **F.** Induction of the p53 protein by licofelone in p53-null RKO colon cancer cells with re-introduced wt p53, but not in cells expressing p53(33/37) as assessed by Western blotting. **G.** Licofelone disrupted the interaction between MDM2 and wtp53 but not p53(33/37) mutant p53 in RKO cells as examined by co-immunoprecipitation. **H.** RITA, PpIX, and licofelone, but not nutlin, disrupted p53/MDMX interaction in HCT 116 cells assessed as in **G**.

To evaluate the role of the structural element which we found to be critical for the allosteric activation of p53, we tested if alanine substitutions of S33 and S37 will prevent the effect of licofelone, using isogenic human colon cancer cells expressing either wtp53 or p53(33/37). Licofelone blocked the interaction between p53 and MDM2 but did not affect the p53(33/37)/MDM2 complex as shown in the immunoprecipitation assay (Figure 7F), suggesting that residues 33 and 37 are important for licofelone-mediated effect. In line with these data, treatment with licofelone induced wtp53, but not p53(33/37) accumulation in RKO cancer cells (Figure 7G).

Finally, we tested if licofelone can prevent p53/MDMX interaction. As evident from the pull-down assay (Figure 7H), RITA, licofelone and PpIX, but not nutlin, blocked the p53/MDMX interaction in HCT116 cells. This indicates that targeting the identified structural element in p53 leads to its activation via a simultaneous inhibition of MDM2 and MDMX binding.

In conclusion, the biological validation of licofelone, a known anti-inflammatory compound, confirmed our prediction that it acts on p53/MDM2 interaction via a newly discovered allosteric mechanism.

## Discussion

Reconstitution of the p53 tumor suppressor leads to the preferential suppression of highly malignant lesions (Feldser et al., 2010 Junttila et al., 2010). These data provide strong support for the studies aimed at reactivation of p53 function. Several compounds targeting the p53/MDM2 interaction via steric hindrance are currently undergoing clinical trials (Hoe et al., 2014). The high attrition rate of new candidate drugs in clinical trials demands the identification of novel compounds with a distinct mode of action and elucidation of mechanisms regulating p53/MDM binding for the efficient implementation of p53-targeting treatments into clinical practice.

Small molecule RITA activates wild-type p53 and inhibits p53/MDM2 interaction, however, it is unique among known MDM2 inhibitors because it binds p53, but not MDM2 (Issaeva et al., 2004). Even though RITA has been reported to display p53-independent effects through the induction of the DNA damage response (Wanzel et al., 2016), it is a valuable tool to explore the mechanism of wild-type p53 reactivation. Thus, here, we addressed the question of how exactly the binding of RITA to p53 affects the p53/MDM2 complex.

In this study we identified the site in the p53 N-terminus targeted by RITA and demonstrated that RITA increases the content of secondary structure in Np53, using CD spectroscopy and IM-MS. Modulation of protein conformation by a weak binding ligand has previously been shown by IM-MS (Harvey et al., 2012). Interestingly, IM-MS can detect the changes in p53 conformation induced by single point mutations in the p53 core domain (Jurneczko et al., 2013) analogous to the detection of the structural changes in Np53 induced by RITA as detected by IM-MS.

Molecular dynamic simulations suggest that residues 32-37, coinciding with RITA-binding site, might be involved in the stabilization of a conformational state in which MDM2-contacting residues F19, W23, and L26 of p53 are tacked inside the molecule (Espinoza-Fonesca, 2009). In line with this study, the tryptophan fluorescence assay demonstrated that W23 is shielded from the solvent (Kar et al., 2002). Since X-ray structure of the MDM2 in complex with short p53 peptide (1YCQ.pdb) suggests that residues F19, W23 and L26 should be facing out to bind MDM2 as shown in Figure 3C (Kussie et al., 1996), it follows that the binding of p53 to MDM2 requires conformational changes. Importantly, a recent study revealed that segments 23-31 and 31-53 of the p53 N-terminus are involved in long-range interactions and can affect p53’s structural flexibility upon MDM2 binding or phosphorylation of residues S33, S46 and T81. In particular, the authors identified nonrandom structural fluctuations at 31-53 segment, which are affected by MDM2 binding. Moreover, the phosphorylation of S33, S46, and T81 induce both local and remote structural changes, which are propagated to the MDM2 binding site in p53 (Lum et al., 2012; Huang et al., 2009). These data provide experimental evidence supporting our idea that restricting conformational mobility of segment involving residues 33-37 might serve to prevent p53/MDM2 interaction.

Figure 6E illustrates our model demonstrating that p53 exists in a range of conformational states, which are present in cells in a dynamic equilibrium. Binding to MDM2 induces F19, W23, and L26 to be exposed and to fit into the p53-binding cleft of MDM2, causing the equilibrium to shift in favour of this conformation. Interaction of RITA with SPLPS stabilizes the alternative conformation, in which MDM2-contacting residues are trapped inside. In this way, the binding of RITA to p53 shifts the balance towards the p53 conformer with low affinity to MDM2.

Further, we found that PpIX, which we have shown to bind to p53 and to disrupt p53/MDM2 interaction, also requires serine 33 and 37 for its functioning as a p53 activator.

Allosteric mechanism of RITA and PpIX action is an unexpected turn in the development of inhibitors of p53/MDM2 interaction, which is mostly focused on steric hindrance mechanism (Hoe et al., 2014).

Intriguingly, RITA-contact residues S33/P34 are involved in the conformation change elicited by Pin1 prolyl-isomerase, an endogenous allosteric modulator for p53, leading to the dissociation of p53 from MDM2 (Mantovani et al., 2007). It is thus possible that the conformational change induced by RITA is similar to the one triggered by Pin1 in cells. Further support for this idea comes from the observations that both RITA and Pin1 cause the release of p53 from its inhibitor iASPP (Issaeva et al., 2004, Mantovani et al., 2007) and that Pin1 contributes to p53-mediated apoptosis induced by RITA (Sorrentino et al., 2013).

One could question that the interaction between RITA and p53 might be too weak to affect the affinity of p53 to MDM2. However, more recent MD simulation shows that hydrogen bonds between residues Phe19-Gln72 and Leu26-Val93 of p53 and MDM2 respectively are unstable and that the total non-bonded interaction energy between p53 and MDM2 is – 162 kcal/mol (Liu and Yan, 2017). This model highlights the importance of residue Trp23 in stabilizing the interaction between p53 and MDM2. Once Trp23 is shielded from the solution, as indicated by our model, it makes the interaction between p53 and MDM2 impossible.

Next, we found that the interaction of p53 with another inhibitor, MDMX, is hindered by an allosteric mechanism. Our study shows that RITA, PpIX, and licofelon inhibit both MDM2 and MDMX binding to p53. This is in line with our data that, in contrast to nutlin, RITA is highly efficient in killing cancer cells with high expression of MDMX (Spinnler et al., 2011).

Our allosteric model of p53/MDM2 interaction allowed us to identify a potent functional analog of RITA, licofelone, which acts similar to RITA.

Anti-inflammatory compound licofelone, a 5-LOX/COX inhibitor has fewer side effects than conventional drugs (Kulkarni et al., 2008). Licofelone has been shown to suppress the growth of cancer cells *in vitro* and *in vivo*, which is attributed to 5-LOX/COX inhibition (Mohammed et al., 2011; Tavolari et al., 2008). Using chemoinformatics, we predicted a new activity of licofelone, i.e., prevention of p53/MDM2 interaction. We confirmed this prediction, showing that licofelone inhibits p53/MDM2 binding and that residues S33/S37 are required for this effect of licofelone. Further, licofelone induced p53 and its target genes and suppressed the growth of cancer cells in a p53-dependent manner.

Prediction of new applications for existing drugs using computational methods has provided fascinating insights into the previously unknown pharmacology of these drugs (Keiser et al., 2009; Murtazalietva et al. 2017). Drug repositioning is an accelerated route of drug discovery, due to the established clinical and pharmacokinetic data. Identification of licofelone as a new inhibitor of p53/MDM2 and MDMX interaction might in future initiate its application for cancer treatment.

Our data establish the allosteric mechanism of inhibition of p53/MDM2 and p53/MDMX interaction by small molecules as a viable strategy for drug discovery. Next, the identified structural elements may provide the basis for the generation of new allosteric activators of p53, which might be valuable additions to the targeted therapeutic pharmacopeia.

## Materials and methods

### In situ proximity ligation assay (PLA)

*In situ* PLA was performed according to the Duolink (Olink biosciences) protocol with modifications (see **Supplemental Experimental procedures** for details).

### Binding assays with [14C]-RITA

For a small molecule-band shift assay purified proteins (20 μM) or 80 μg of total protein from cell lysates and [^14^C]-RITA (40 μM) were incubated in buffer B (50 mM Hepes, pH 7.0, 150 mM NaCl, 35% glycerol) at 37°C 30 min and separated in standard TBE or gradient native gels. Gels were stained to visualize proteins and radioactivity was measured using Phosphoimager Amersham Biosciences.

SDS-PAGE separation of RITA/protein complexes was performed in 10% gel after brief heating at 90°C of lysates in loading buffer. Proteins were depleted from cell lysates using anti-p53 DO-1 or anti-actin antibody (AC-15, Sigma), immobilized on protein A-conjugated DynaBeads (Invitrogen).

Co-immunoprecipitation of p53/MDM2 or p53/MDMX was performed as described previously (Issaeva et al., 2004). MDM2 in precipitates from mouse tumor cells MCIM SS cells expressing wtp53 (Magnusson et al., 1998) was detected by the 4B2 antibody, a gift from Dr. S. Lain. MDMX antibody was from Bethyl laboratories.

### Scintillation Proximity Assay

SPA PVT Protein A beads (500μl/sample) were incubated for 2h with anti-GST antibodies (1:100). 0.1 μg/μl of studied protein in SPA buffer (GST, Np53, Np53(33/37) was added to GST-coated SPA beads. 10 μl [^14^C]-RITA (52 μCi) diluted 4 times in SPA buffer were added to protein samples (1.3 μCi). Unlabelled RITA was used as a cold substrate. SPA buffer was added to the final volume of 100 μl. Complexes were incubated for 1h at 37°C and luminescence released by the [^14^C]-RITA-excited beads were measured in a standard microplate reader.

### Circular dichroism spectroscopy (CD)

Proteins 50 μM were incubated with RITA (reconstituted in 100% isopropyl alcohol, IPA) or with the same amount of IPA as a blank at a 1:2 molar ratio in 25 mM ammonium acetate at 37°C for 20 min. This results in a final concentration of IPA in each case of 5%.

All CD spectra were acquired using a JASCO instrument. 0.1 cm Hellma^®^ cuvettes were used and measurements were performed in the far-UV region 260 – 195 nm at 21°C. CD spectra were recorded with a 1 nm spectral bandwidth, 0.5 nm step size with scanning speed 200 nm/min. The spectra were recorded 5 times, and the data are representative of at least three independent experiments.

### Mass Spectrometry and Ion Mobility Mass Spectrometry (IM-MS)

Mass spectrometry and IM-MS were made on an in-house modified quadrupole time-of-flight mass spectrometer (Waters, Manchester, UK) containing a copper coated drift cell of length 5cm. The instrument, its operation and its use in previous studies on p53 have been described elsewhere (McCullough et al., 2008; Jurneczko et al., 2013). Np53 was prepared at a concentration 50 μM in 50 mM Ammonium Acetate. Protein was incubated with RITA at a 1:2 molar ratio at 37°C for 30 minutes before analysis. 5% isopropyl alcohol was added to solubilize the ligand in aqueous solution, consistent with CD spectroscopy data. In all cases three repeats were taken, each on different days (For details see **Supplemental Experimental procedures**).

### Molecular Modelling

Homology model of p53 was developed using the Rosetta server (Kim et al., 2004, 2005; Rohl et al., 2004; Chivian et al., 2006). Generated models were validated and fitted to the cryo-EM data (Okorokov et al., 2006). Domain fitting into the 3D map of p53 was performed automatically using UCSF Chimera package from the Resource for Biocomputing, Visualization, and Informatics at the University of California, San Francisco (supported by NIH P41 RR-01081), (www.cgl.ucsf.edu/chimera/) and further refined by UROX (http://mem.ibs.fr/UROX/). (For details see **Supplemental Experimental procedures**).

### Yeast-based reporter assay

The yeast-based functional assay was conducted as previously described ^3^. Briefly, the p53-dependent yeast reporter strain yLFM-PUMA containing the luciferase cDNA cloned at the *ADE2* locus and expressed under the control of PUMA promoter (Inga et al., 2002) was transfected with pTSG-p53 (Resnick and Inga 2003), pRB-MDM2 (generously provided by Dr. R. Brachmann, Univ. of California, Irvine, CA, USA), or pTSG-p53 S33/37 mutant and selected on double drop-out media for TRP1 and HIS3. Luciferase activity was measured 16 hrs after the shift to galactose-containing media and the addition of 1 μM RITA, PpIX or 10 μM nutlin (Alexis Biochemicals), or DMSO. Presented are average relative light units and the standard errors obtained from three independent experiments each containing five biological repeats.

## Supporting information

## Supplemental information

Supplemental information includes Supplemental Experimental Procedures, Supplemental References, four supplemental figures and two tables.

## Acknowledgments

This work was supported by grants to G.S. from the Swedish Research Council, the Swedish Cancer Society, and Ragnar Söderberg Foundation. JZP would like to acknowledge the grant from Karolinska Institute, Stockholms Läns Ladsting, the Strategic Research Program in Cancer and Åke Wibergs Stiftelse. O.T. and V.P would like to acknowledge the Russian State Academies of Sciences Fundamental Research Program for 2013-2020. Special thanks are addressed to Prof. Sir David Lane for helpful discussions, suggestions and encouraging feedback. We thank Prof. Sonia Lain for providing ARN8 and T22 cells and helpful discussions. We would like to thank Dr. Margareta Wilhelm for providing a Ras, cMyc transformed mouse embryonic fibroblast and helpful discussions. The authors are grateful to Yari Ciribilli, Bartosz Ferens, Dr. Anna Kostecka and Dr. Alicja Sznarkowska for technical assistance and helpful discussions. We are greatly indebted to Protein Science Facility Karolinska Institutet for protein purification and to all our colleagues who shared with us their reagents and cell lines.

## Authors contribution

Conceptualization: G.S., J.Z-P; A.L.O. Methodology and data analysis: G.S., J.Z-P, A.L.O, V.V.G., N.I., P.E.B., A.I., LG.L, A.K., V.P., M.P. Investigation: J.Z-P., V.V.G., M.B., A.V., K.R., N.I., V.A., E.R.D., E.H., C.S., O.T., S.L.; Writing draft: J.Z-P., A.L.O, V.V.G, V.P., A.I., LG.L., O.T., P.E.B, G.S.; Writing review and editing: J.Z-P.; Supervision: G.S. and J.Z-P.

## References

Andreotti, V. p53 transactivation and the impact of mutations, cofactors and small molecules using a simplified yeast-based screening system. PLoS One. 6(6), e20643 (2011).

Castell, A. et al. A selective high-affinity MYC-binding compound inhibits MYC: MAX interaction and MYC-dependent tumor cell proliferation. Sci Rep. 8(1), 10064 (2018).

Chivian, D., Baker, D. Homology modeling using parametric alignment ensemble generation with consensus and energy-based model selection. Nucleic Acids Res., 34, e112 (2006).

Dickinson, E. R. et al. The use of ion mobility mass spectrometry to probe modulation of the structure of p53 and of MDM2 by small molecule inhibitors. Front Mol Biosci, 2, 39 (2015).

Dudgeon, C. et al. Tumor susceptibility and apoptosis defect in a mouse strain expressing a human p53 transgene. Cancer Res. 15:66(6), 2928–2936 (2006).

Enge, M. et al. MDM2-dependent downregulation of p21 and hnRNP K provides a switch between apoptosis and growth arrest induced by pharmacologically activated p53. Cancer Cell 15(3), 171–183 (2009).

Espinoza-Fonesca, L.M. Leucine-rich hydrophobic clusters promote folding of the Nterminus of the intrinsically disordered transactivation domain of p53. FEBS Lett. 583(3), 556–560 (2009).

Feldser, D.M. et al. Stage-specific sensitivity to p53 restoration during lung cancer progression. Nature. 468(7323), 572–5 (2010).

Filimonov, D.A. et al. Prediction of the biological activity spectra of organic compounds using the PASS online web resource. Chem. Heterocycl. Compd. 50(3), 444–457 (2014).

Grinkevich, V., Issaeva, N., Hossain, S., Pramanik, A., Selivanova, G. Reply to ‘NMR indicates that the small molecule RITA does not block p53-MDM2 binding in vitro’. Nat Med. 11, 1136–1137 (2005).

Grinkevich, V. et al. Ablation of key oncogenic pathways by RITA-reactivated p53 is required for efficient apoptosis. Cancer Cell. 15(5), 441–453 (2009).

Harvey, S. R. et al. Small-Molecule Inhibition of c-MYC:MAX Leucine Zipper formation Is Revealed by Ion Mobility Mass Spectrometry. J. Am. Chem. Soc. 134, 19384–19392 (2012).

Hoe, K. K., Verma, C. S, Lane, D. P. Drugging the p53 pathway: understanding the route to clinical efficacy. Nat Rev Drug Discovery, 13, 217–236 (2014).

Huang, F. et al. Multiple conformations of full-length p53 detected with single-molecule fluorescence resonance energy transfer. Proc Natl Acad Sci U S A. 106:49, 20758–20763 (2009).

Inga, A., Storici, F., Darden, T.A., Resnick, M.A. Differential transactivation by the p53 transcription factor is highly dependent on p53 level and promoter target sequence. Mol Cell Biol. 22(24), 8612–25 (2002).

Issaeva, N. et al. Small molecule RITA binds to p53, blocks p53-HDM-2 interaction and activates p53 function in tumors. Nat. Med. 10, 1321–1328 (2004).

Junttila, M. R. et al. Selective activation of p53-mediated tumour suppression in high-grade tumours. Nature. 468(7323), 567–71 (2010).

Jurneczko, E. et al. Probing the Conformational Diversity of Cancer-Associated Mutations in p53 with Ion-Mobility Mass Spectrometry. Angew. Chem. Int. Ed. 52, 4370–4374 (2013).

Kar, S. et al. Effect of Phosphorylation on the Structure and Fold of Transactivation Domain of p53. J. Biol. Chem. 277, 15579 – 15585 (2002).

Keiser, M. J. et al. Predicting new molecular targets for known drugs. Nature. 462(7270), 175–81 (2009).

Kim, D. E., Chivian, D., Baker, D. Protein structure prediction and analysis using the Robetta server. Nucleic Acids Res. 32, W526 – W531 (2004).

Kim, D. E., Chivian, D., Malmström, L., David Baker, D. Automated prediction of domain boundaries in CASP6 targets using Ginzu and RosettaDOM. Proteins: Struct. Funct., Bioinf. 61, 193–200 (2005).

Koehler, M. F. et al. Albumin affinity tags increase peptide half-life in vivo. Bioorg Med Chem Lett. 12(20), 2883–2886 (2002).

Krajewski, M., Ozdowy, P., D’Silva, L., Rothweiler, U., Holak, T. A. NMR indicates that the small molecule RITA does not block p53-MDM2 binding in vitro. Nat. Med. 11(11), 1135– 1136 (2005).

Kulkarni, S. K., Singh, V. P. Licofelone: the answer to unmet needs in osteoarthritis therapy? Curr Rheumatol Rep. 10(1), 43–8 (2008).

Kussie, P. H. et al. Structure of the MDM2 oncoprotein bound to the p53 tumor suppressor transactivation domain. Science. 274(5289), 948–953 (1996).

Lagunin, A., Filimonov, D. A., Poroikov, V. V. Multi-targeted natural products evaluation based on biological activity prediction with PASS. Cur. Phar. Des. 16(15), 1703–1717 (2010).

Liu, S. X., Yan, S. W. Mechanism of competition between Nutlin 3 and p53 for binding with MDM2. Chin. Phys. Let. 34(11), 118701 (2017).

Lowry, D. F., Hausrath, A. C., Daughdrill, G. W. A robust approach for analyzing a heterogenous structural ensemble. Proteins: Struct., Funct., Bioinf. 73, 918–928 (2008).

Mohammed, A. et al. Chemoprevention of colon and small intestinal tumorigenesis in APC(Min/+) mice by licofelone, a novel dual 5-LOX/COX inhibitor: potential implications for human colon cancer prevention. Cancer Prev Res (Phila). 4(12), 2015–26 (2011).

MacroModel, version 9.6; Schrödinger, LLC: New York, NY, (2008).

Magnusson, K. P., Satalino, R., Qian, W., Klein, G., Wiman, K. G. Is conversion of solid into more anoxic ascites tumors associated with p53 inactivation? Oncogene 17, 2333–2337 (1998).

Mantovani, F. et al. The prolyl isomerase Pin1 orchestrates p53 acetylation and dissociation from the apoptosis inhibitor iASPP. Nat Struct Mol Biol. 14, 9212–9220 (2007).

McCullough, B. J. et al. Development of an Ion Mobility Quadropole Time of Flight Mass Spectrometer. Anal. Chem. 80, 6336–6344 (2008).

Murtazalieva, K. A., Druzhilovskiy, D. S., Goel, R. K., Sastry, G. N., Poroikov, V. V. How good are publicly available web services that predict bioactivity profiles for drug repurposing? SAR QSAR in Environ Res. 28(10), 843–862 (2017).

Okorokov, A.L., Sherman, M.B., Plisson, C., Grinkevich, V., Sigmundsson, K., Selivanova, G., Milner, J., Orlova, E.V. The structure of p53 tumour suppressor protein reveals the basis for its functional plasticity. EMBO J. 25(21), 5191–5200 (2006).

Resnick, M.A. & Inga, A. Functional mutants of the sequence-specific transcription factor p53 and implications for master genes of diversity. Proc Natl Acad Sci U S A. 100(17), 9934–9 (2003).

Rohl, C. A., Strauss, C. E. M., Chivian, D., Baker, D. Modeling structurally variable regions in homologous proteins with rosetta. Proteins 55, 656–677 (2004).

Sorrentino, G. et al. The prolyl-isomerase Pin1 activates the mitochondrial death program of p53. Cell Death Differ. 20(2), 198–208 (2013).

Söderberg, O. et al. Direct observation of individual endogenous protein complexes in situ by proximity ligation. Nat. Methods. 3, 995–1000 (2006).

Sur, S. et al. A panel of isogenic human cancer cells suggests a therapeutic approach for cancers with inactivated p53. Proc Natl Acad Sci U S A. 106(10), 3964–3996 (2009).

Shangary, S. et al. Reactivation of p53 by a specific MDM2 antagonist (MI-43) leads to p21-mediated cell cycle arrest and selective cell death in colon cancer. Mol Cancer Ther. 7, 1533–1542 (2008).

Sznarkowska, A. et al. Targeting of p53 and its homolog p73 by protoporphyrin IX. FEBS Lett. 585(1), 255–260 (2011).

Tavolari, S. et al. Licofelone, a dual COX/5-LOX inhibitor, induces apoptosis in HCA-7 colon cancer cells through the mitochondrial pathway independently from its ability to affect the arachidonic acid cascade. Carcinogenesis 29(2), 371–80 (2008).

Toledo, F., Wahl, G. M. MDM2 and MDM4: p53 regulators as targets in anticancer therapy. Int J Biochem Cell Biol. 39, 1476–1482 (2007).

Vousden, K. H., Prives, C. Blinded by the Light: The Growing Complexity of p53. Cell. 137(3), 413–31 (2009).

Uversky, V. N. p53 Proteoforms and Intrinsic Disorder: An Illustration of the Protein Structure–Function Continuum Concept. Int J Mol Sci. 17(11), 1874 (2016).

Yang, Y. et al. Small molecule inhibitors of HDM2 ubiquitin ligase activity stabilize and activate p53 in cells. Cancer Cell. 7(6), 547–559 (2005).

Zawacka-Pankau, J. et al. Protoporphyrin IX interacts with wild-type p53 protein in vitro and induces cell death of human colon cancer cells in a p53-dependent and independent manner. J Biol Chem. 282(4), 2466–2472 (2007).

Wanzel, M. et al. CRISPR-Cas9-based target validation for p53-reactivating model compounds. Nat Chem Biol. 12(1), 22–8 (2016).

