## Supplementary material for "Novel allosteric mechanism of p53 activation by small molecules for targeted anticancer therapy"

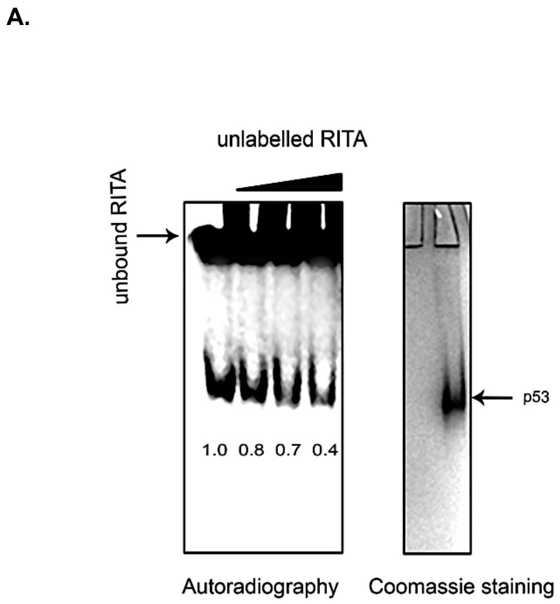
A.
unlabelled RITA
unbound RITA
1.0 0.8 0.7 0.4
p53
Autoradiography
Coomassie staining

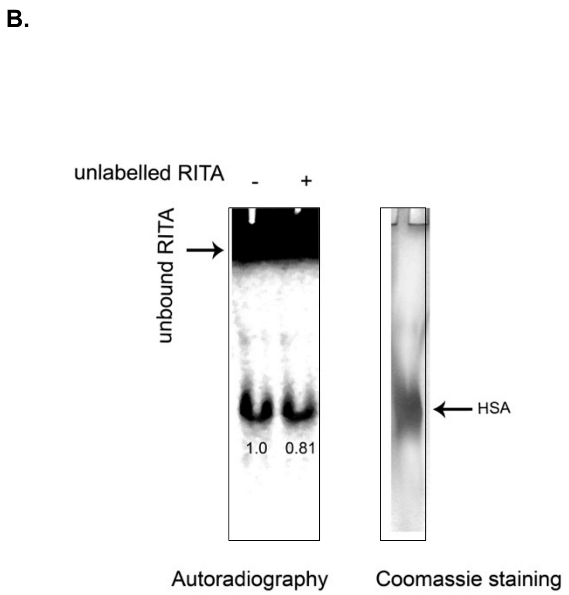
B.
unlabelled RITA
-
+
unbound RITA
1.0 0.81
HSA
Autoradiography
Coomassie staining

Zawacka-Pankau_FigS2

**A.**

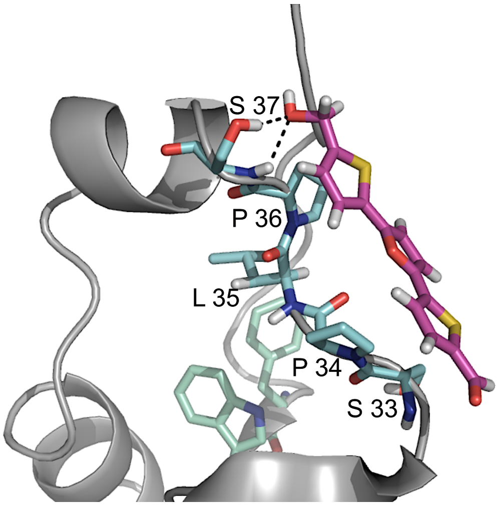


**B.**

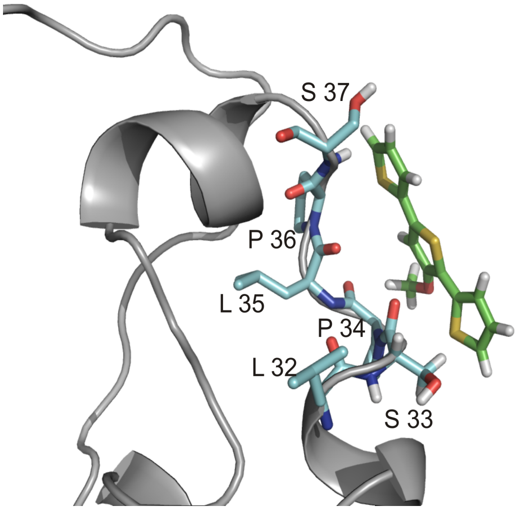

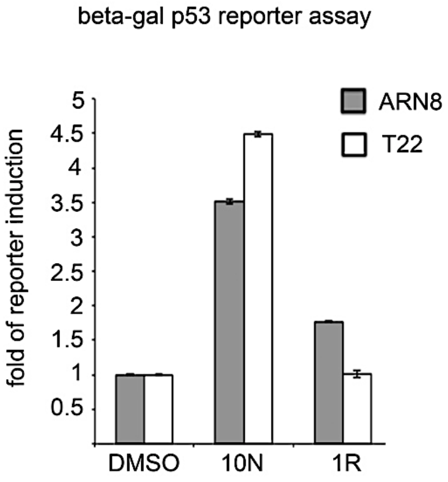
beta-gal p53 reporter assay
fold of reporter induction
5
4.5
4
3.5
3
2.5
2
1.5
1
0.5
ARN8
T22
DMSO
10N
1R

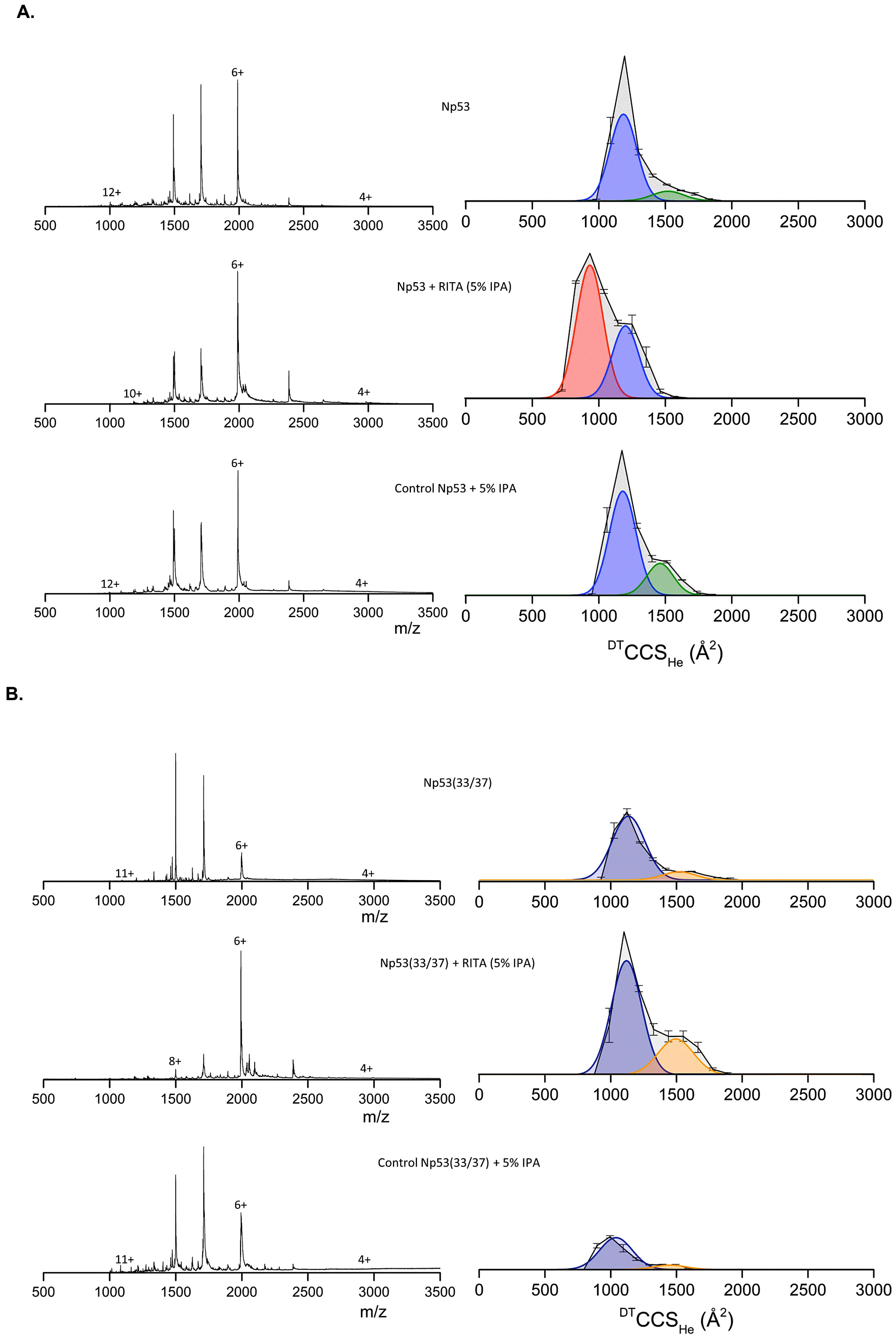
A.
Np53
Np53 + RITA (5% IPA)
Control Np53 + 5% IPA
12+
6+
4+
10+
m/z
$^{DT}CCS_{He}$ (Å$^2$)
B.
Np53(33/37)
Np53(33/37) + RITA (5% IPA)
Control Np53(33/37) + 5% IPA
11+
8+

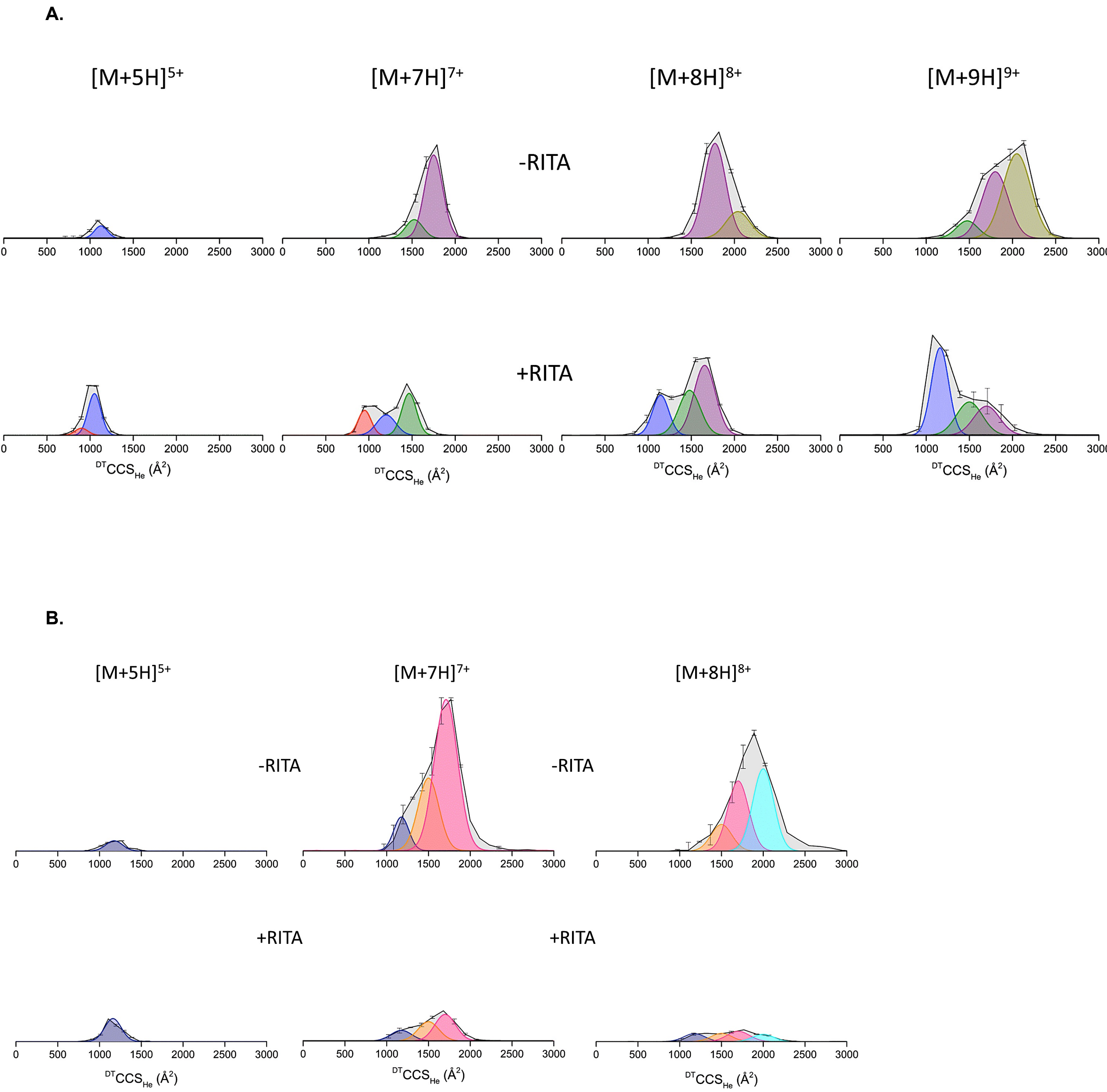
A.
[M+5H]5+
[M+7H]7+
[M+8H]8+
[M+9H]9+
-RITA
+RITA
DTCCSHe (Å2)
B.
[M+5H]5+
[M+7H]7+
[M+8H]8+
-RITA
-RITA
+RITA
+RITA
DTCCSHe (Å2)

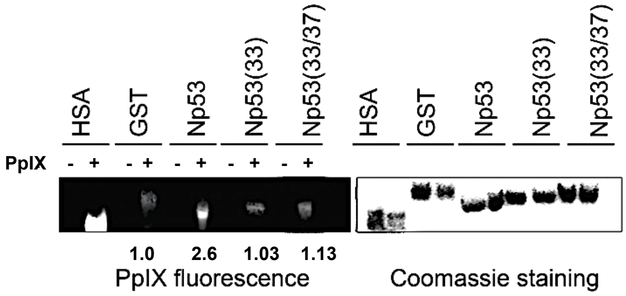
HSA
GST
Np53
Np53(33)
Np53(33/37)
HSA
GST
Np53
Np53(33)
Np53(33/37)
PpIX - + - + - + - + - +
1.0 2.6 1.03 1.13
PpIX fluorescence
Coomassie staining

A.

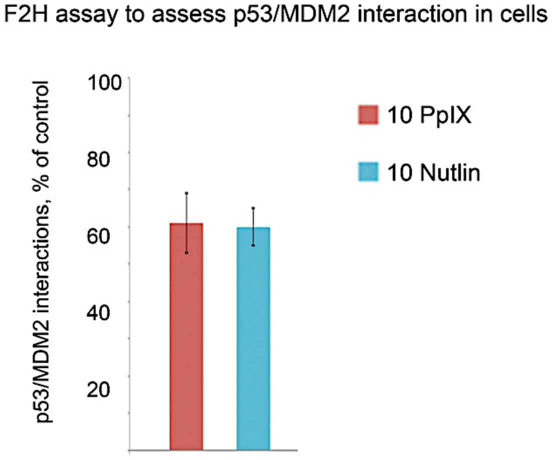
F2H assay to assess p53/MDM2 interaction in cells
p53/MDM2 interactions, % of control
100
80
60
40
20
10 PpIX
10 Nutlin

B.

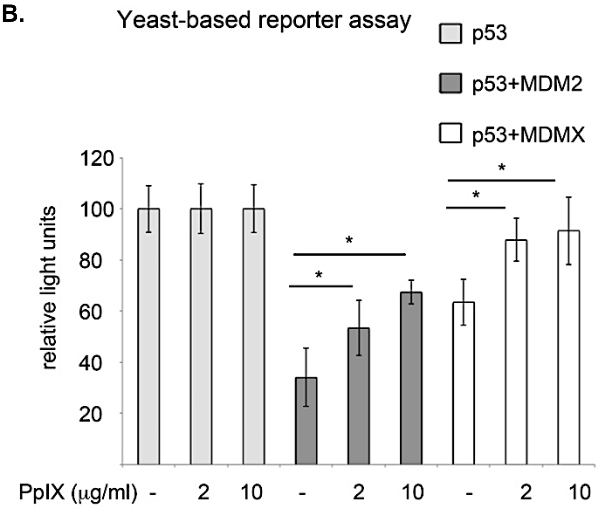
Yeast-based reporter assay
p53
p53+MDM2
p53+MDMX
relative light units
120
100
80
60
40
20
*
*
*
*
PpIX (μg/ml) - 2 10 - 2 10 - 2 10

**Structural analogs of RITA and their biological activities**

| Lp. | ID | Structure | Active/inactive |
|---|---|---|---|
| 1 | RITA | 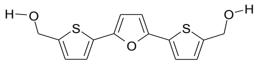 | Active (RITA) |
| 2 | LCTA-2081 | 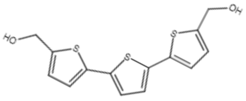 | Active |
| 3 | NSC672170 | 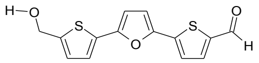 | Active (lower than RITA) |
| 4 | NSC650973 | 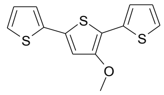 | Inactive |
| 5 | NSC629035 | 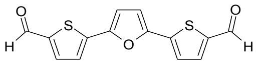 | Inactive |
| 6 | NSC613590 | 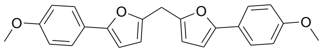 | Inactive |
| 7 | NSC691803 | 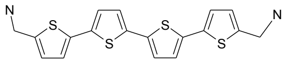 | Inactive |
| 8 | NSC647123 | 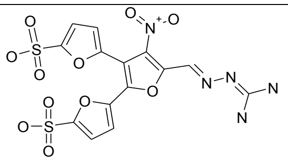 | Inactive |
| 9 | NSC657767 | 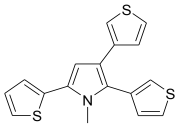 | Inactive |
| 10 | NSC661061 | 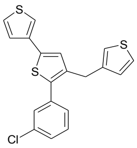 | Inactive |
| 11 | NSC694946 | 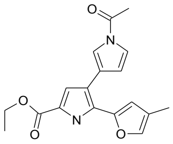 | Inactive |
| 12 | NSC116644 | 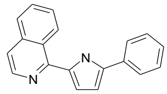 | Inactive |

**Functional analogs of RITA identified using chemoinformatics**

| | DRUG ID | DRUG NAME | STRUCTURE | TANIMOTO COEFFICIENT |
|---|---|---|---|---|
| 1 | DCL000302 | LICOFELONE | 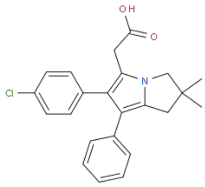 | 0.730445 |
| 2 | DCL000651 | TAK-442 | 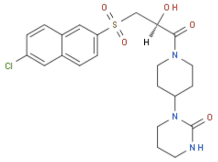 | 0.718800 |
| 3 | DCL000306 | BENZBROMARONE | 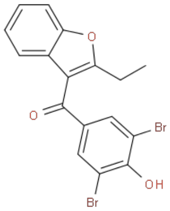 | 0.655255 |
| 4 | DCL000444 | AVE0010 | 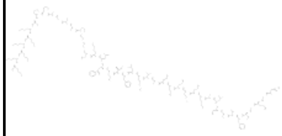 | 0.654472 |
| 5 | DAP000792 | RALOXIFENE | 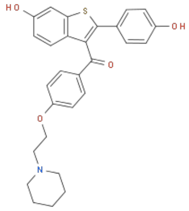 | 0.653526 |
| 6 | DAP000115 | ADAPALENE | 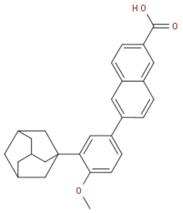 | 0.652967 |
| 7 | DAP000089 | PROPRANOLOL | 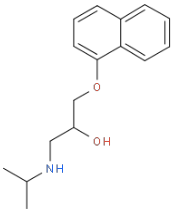 | 0.596243 |

| | DRUG ID | DRUG NAME | STRUCTURE | TANIMOTO COEFFICIENT |
|---|---|---|---|---|
| 8 | DCL000615 | R7204 | 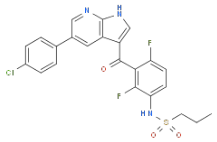 | 0.589953 |
| 9 | DAP000554 | FLUVASTATIN | 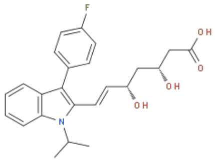 | 0.588334 |
| 10 | DCL000100 | DAP-81 | 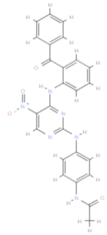 | 0.580234 |
| 11 | DAP000968 | NAPROXEN | 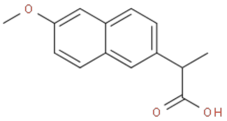 | 0.580234 |
| 12 | DCL000835 | HCV-796 | 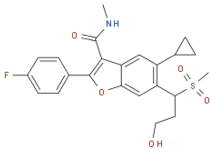 | 0.554923 |
| 13 | DAP000735 | NABUMETONE | 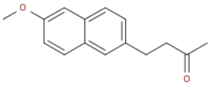 | 0.545487 |
